# PIEZO1-dependent erythrocyte dehydration as the mechanism for selection of an allele protecting from severe malaria

**DOI:** 10.1101/2022.03.31.486604

**Authors:** Svetlana Glushakova, Ludmila Bezrukov, Hang Waters, Yuto Kegawa, Paul S. Blank, Joshua Zimmerberg

## Abstract

PIEZO1 is a cation specific mechanoreceptor channel implicated in red blood cell (RBC) volume homeostasis. Several PIEZO1 gain of function (GoF) variants demonstrate delayed channel inactivation and can cause hereditary xerocytosis (HX), a disease characterized by hemolytic anemia, RBC dehydration, and shape distortion. The milder PIEZO1_E756del_ GoF variant, prevalent in populations of African descent, protects carriers from severe malaria caused by *Plasmodium falciparum* and ameliorate disease in a rodent malaria model. To explore the mechanism of this malaria protection, *P. falciparum* infection of human PIEZO1_E756del_ RBC was analyzed in shear-stressed and static cultures with and without Yoda1, a PIEZO1 agonist. RBC dehydration was a common pathophysiological factor affecting parasite replication in both culture conditions. PIEZO1 channel opening by either Yoda1 or shear stress produced dehydration-dependent cell hemolysis, inhibiting *P. falciparum* infection. Since the physiological activator of PIEZO1 in circulating RBC is shear stress, we propose that shear stress-induced dehydration, disproportionally affecting RBC of GoF PIEZO1 _E756del_ carriers, makes erythrocytes less habitable for *P. falciparum* to the point of hemolysis, and thus ameliorates malaria in GoF PIEZO1_E756del_ carriers. More generally, RBC dehydration processes may be a pathway for protection from the severe form of malaria common to several hematological disorders, including sickle cell trait.

**Key points:**

- PIEZO1_E756del_ activation in African American donor RBC provokes dehydration-dependent cell hemolysis, impairing *P. falciparum* replication.
- RBC dehydration could be a malaria ameliorating factor in several known RBC hematological disorders, including sickle cell trait.

## Introduction

Reflecting an evolutionary pressure of malaria parasites, malaria-endemic regions sustain high incidences of hematological disorders targeting human erythrocytes, the host cells for malaria parasite replication.^1,2^ The gain of function (GoF) mutation in a human mechanosensitive cation channel, PIEZO1_E756del_, is present in high frequency amongst populations of African descent.^3^ Some GoF PIEZO1 mutations are associated with hereditary xerocytosis (HX), a hemolytic anemia characterized by distorted and dehydrated RBC.^4-6^ When infected with *P. berghei*, mice engineered with the PIEZO1_E756del_ mutation had lower parasitemia and mortality from cerebral malaria than control mice.^3^ African malaria patients with heterozygous PIEZO1_E756del_^7^ or homozygous PIEZO1_E756del_^8^ have a lower probability of severe malaria. The link between the PIEZO1_E756del_ mutation and malaria is the first example of an ion channel dysfunction ameliorating malaria.

The cellular mechanism explaining the association between PIEZO1_E756del_ variant and malaria is still unknown. Abnormal exposure of knobs on the surface of infected human RBC with PIEZO1 E756del (iRBC_E756del_) may preferentially eliminate iRBC by the spleen *in vivo*^7,9,10^ or parasite replication may be defective in RBC_E756del_.^3,11^ Limited knowledge of both the physiological role of PIEZO1 in RBC and the effect of PIEZO1_E756del_ variant on RBC physiology confounds studies on cellular mechanisms of malaria amelioration in carriers of this mutation. PIEZO1 channel properties are studied using patch-clamp electrophysiology, directly recording ion currents.^12,13^ PIEZO1, which is a non-selective, membrane cation channel, is normally activated, i.e., opened, upon application of a physical strain stressing the cell membrane, allowing an ionic current to pass.^12,14^ Both HX-associated and E756del GoF mutations in PIEZO1 demonstrate delayed channel inactivation (closing of the channel), albeit less severe in the latter case.^3,5,15^ The PIEZO1 agonist Yoda1 activates PIEZO1 channel like mechanical activation^16^ making Yoda1 useful for experiments in cell suspensions. Both chemical and mechanical channel activations lead to an influx of Ca^2+^ into RBC activating Ca^2+^-dependent potassium channels (KCNN4 also known as the Gardos channel), subsequent efflux of intracellular K^+^ through KCNN4, and resulting in a quick reduction in cell volume.^17,18^ RBC stretching during passage through a narrow microfluidic system (mimicking capillary vessels) and in mice capillaries *in vivo*, activates PIEZO1 and increases the concentration of RBC intracellular free calcium ([Ca^2+^]_free_).^19^ The high Ca^2+^ gradient across the erythrocyte membrane (∼ 2 mM in human plasma vs. ∼ 60 nM inside a RBC) facilitates a fast [Ca^2+^]_free_ increase during the time of the channel’s open state while undergoing capillary transit.^20^ Because of delayed channel inactivation in GoF PIEZO1^5,15^, a more profound cell dehydration should result in these variants, as rapid Ca^2+^ entry would overwhelm the Ca^2+^ pump and a loss of intracellular KCl, described above, would ensue.^17,20^ Indeed, GoF PIEZO1 erythrocytes are more dehydrated and have a lower osmotic fragility^3^ even under *in vitro* static conditions where the channel presumably is inactive^17^ due to an absence of shear stress-inducing channel activation, as described for other cell types.^21-23^

Here we explore putative cellular mechanisms and external factors contributing to protection from severe malaria for PIEZO1_E756del_ carriers. A quantitative optical microscopy analysis of parasite replication in static culture^24-26^ was used to assess all stages of the parasite intraerythrocytic replicative cycle. Parasite replication was the same in RBC_E756del_ with inactive PIEZO1 compared to blood from donors without this mutation (RBC_WT_) but was much less efficient in the dehydrated RBC isolated from blood of PIEZO1_E756del_ carriers. Channel activation by a chemical agonist (Yoda1) and physical force (shear stress) *in vitro* strongly inhibited parasite replication in RBC_E756del_ and was associated with cell-dehydration dependent hemolysis. The close pathophysiology of sickle cell and RBC_E756del_ prompts us to propose that the cell dehydration found in the blood of PIEZO1_E756del_ carriers exemplify a mechanism for the selective advantage of alleles that reduce the burden of malaria.

## Materials and methods

### Identification of PIEZO1 GoF carriers among the blood donors

Anonymized blood samples from donors participating in the National Institutes of Health IRB-approved Research Donor Program in Bethesda, MD were used. Blood was collected into BD Vacutainers with acid citrate dextrose additives. Genomic DNA was isolated using QIAamp DNA mini (Qiagen, #51104); genomic PCRs were performed with Platinum PCR Supermix High Fidelity kit (Invitrogen, #12532016) using primer sets to identify three targeted GoF PIEZO1 mutations E756del, A1988V and R2456H, and sickle cell mutation in hemoglobin-Beta (HbS): (forward/reverse primers) CAGGCAGGATGCAGTGAGTG/ GGACATGGCACAGCAGACTG for E756del; CGCTTCTTCCACGACATCC/ GGAAGTTGCCGAGGATGC for A1988V; CAGTGACAAGGTCAGCCCAC/ CGGTGAGCGGTAGAGGAAG for R2456H; AGTCAGGGCAGAGCCATCTA/ GTCTCCACATGCCCAGTTTC for HbS. PCR products were purified using Qiagen PCR purification kit (Qiagen, #28104) and sequenced by Psomagen, Inc. (Rockville, MD). The resulting sequences were analyzed using Sequencher (Gene Code Corporation, MI) and Vector NTI (Invitrogen, CA), and aligned with a WT PIEZO1 genome sequence from NCBI to identify the PIEZO1 GoF carriers.

### Characterization of erythrocyte physical properties

Blood erythrocytes were assayed for osmotic fragility (reflecting cell hydration), as described in Ma et al., 2018, and cell morphology on the day of donation or within 24 h of donation but prior to blood processing. SigmaPlot (Systat Software, Inc.) was used to determine C_50_, the relative tonicity of the buffer resulting in 50% of the maximum RBC hemolysis. Cell morphology was evaluated at 37^°^C by differential interference contrast (DIC) microscopy using an LSM800 Zeiss microscope (63x oil objective, NA1.4) and HybriWell HBW20 chambers (Grace Bio-Labs, Inc.). On average, 500-1200 RBC were inspected for each donor blood sample. Deviations from rounded, biconcave, RBC shape were classified as “distorted morphology”.

### Isolation of dehydrated cells by density gradient

Percoll (Sigma) was used to isolate dehydrated RBC from donor blood. Discontinuous steps made of isotonic Percoll (90% and 80% with density 1.124 g/L and 1.107 g/L correspondingly) supplemented with 2 mM HEPES and 25 µg/ml gentamicin were used for the gradient. The centrifugation conditions were: 23,000 rpm, 30 min,15 ^°^C, SW50.1 rotor, Optima XE-100 ultracentrifuge (Beckman Coulter). Cells from 90% Percoll were called the “dehydrated cell fraction” (DF) and contained RBC more dehydrated than the typical circulating RBC of healthy donors.^27^ Cells, washed from the Percoll, were kept in complete RPMI medium for subsequent use.

### Analysis of intraerythrocytic parasite replicative cycle in cells with inactive PIEZO1

RBC from donor blood were isolated using a 65% isotonic Percoll step density gradient centrifugation (1500 g for 15 min at 30 ^°^C), washed from the Percoll and stored in complete medium prior to experiments. *P. falciparum* strain NF54 expressing two fluorescent proteins (vacuolar membrane protein EXP2-mNeonGreen and soluble vacuolar protein mRuby3)^28^ was used for infection. A traditional cell culture method, as described in Glushakova et al.^29^, was used to assay *P. falciparum* replication in RBC with inactive PIEZO1. Briefly, RBC (1 ml of 2% hct) were maintained at 37 ^°^C in an atmosphere of 5% O_2_, 5% CO_2_ and 90% N_2_ in complete medium (RPMI 1640, 25 mM HEPES, 0.1 mM hypoxanthine, 25 µg/ml gentamicin, 0.5% Albumax II (all reagents from Gibco), and 4.5 mg/ml glucose (Sigma)). Live-cell microscopy (LSM800, Zeiss) was used to evaluate cell morphology and the erythrocyte cycle progression.^24-26, 28-29^ 488 and 561 nm laser light at 0.2 mW and 0.6 mW was used to excite mNeonGreen and mRuby3, respectively. Highly synchronized cultures of *P. falciparum* were prepared according to Glushakova et al.^25^ Parasite egress and invasion were observed using HybriWell HBW20 chambers.^29^ The individual stages of the parasite cycle were evaluated as described previously.^24-26^ The length of the erythrocyte cycle was evaluated based on an egress assay using highly synchronized cultures. To compare cell cycle length in two different cultures, egress was assayed sequentially every second hour starting from 40-42 hours post infection initiation for 10-12 hours using the same cultures. Cumulative egress was calculated, and the results presented as the kinetics of parasite egress. The kinetic curves were created in Sigma Plot software to quantify initiation and progression of parasite egress.

### Activation of PIEZO1 in vitro

Both chemical (PIEZO1 agonist Yoda1 (Tocris Bioscience)) and mechanical (shear stress) activation of PIEZO1 were used. Preliminary experiments showed that i) parasite egress was not inhibited by Yoda1, ii) pretreatment of normal RBC or treatment of a mixture of infected and normal cells during the initiation of infection gave the same result, and iii) cell treatment with Yoda1 beyond 1h at 37 ^°^C did not affect the outcome (data not presented). For parasite replication assays, cells (2% hct, 37 ^°^C) were treated for 1 hour with 5 and 10 uM Yoda1 in complete medium within a 2-hour window of infection initiation.^16^ The drug was removed using three sequential cell washes. The control cultures were treated with DMSO and underwent the same manipulations. The mechanical activation of PIEZO1 in RBC by shear stress was achieved by constant agitation of cells (1-2 × 10^8^ cells/ml) on the platform of a “Belly Button Shaker” (Model BBUAAUV15, Ibi Scientific) at a speed of 14 rpm, the minimal rate of rotation necessary to keep erythrocytes in suspension inside the sterile, 40 mm long by 9 mm wide, rounded bottom tubes.

### Assessment of cell hemolysis

Aliquots of culture medium were assayed for absorption at 540 nm (Microplate Reader Infinite 200 PRO, Nano Quant). The cumulative hemolysis for the entire experiment was calculated and used to compare hemolysis in the different cultures, each with 1-2 × 10^8^ cells/ml.

### Statistical analysis

Data were analyzed using SigmaPlot (Systat Software, Inc., San Jose, CA) and Matlab (The MathWorks, Inc., Natick, MA). RBC hydration and parasitemia data were logit (log_e_ (% Value/(100 - % Value)) and log_e_ transformed prior to statistical evaluation, respectively. Summary data are presented as (mean +/-SEM, n = experiment number) of the untransformed data unless indicated otherwise. For small n where the distributional assumptions required for testing are difficult to validate (t-test, for example), a bootstrap analysis of the difference between two groups, using transformed data, was performed and rejection of the null hypothesis that two groups have the same mean was set at the α = 0.001 level. One factor ANOVA was used for comparisons of 3 groups.

### Data sharing

For original data, please contact Svetlana Glushakova via e-mail at

## Results

### *P. falciparum* infected RBC_E756del_ manifest unusual cytopathology

To assess the effect of intracellular host factors on parasite replication in RBC_E756del_ with presumably inactive PIEZO1, the standard *in vitro* method was employed. Blood from four self-identified African American (AA) donors having GoF PIEZO1 E756del mutation but not carrying PIEZO1 A1988V and R2456H GOF or hemoglobin beta sickle cell mutations were used. The E756del mutation in PIEZO1 induced a mild RBC dehydration (C_50_=41.10+/-1.16%, mean+/-SEM, n=7 from 4 donors). The effect on cell dehydration of the PIEZO1_E756del_ mutation (7 samples vs 53 samples of PIEZO1_WT_ blood) was significant (non-distributional bootstrap analysis; see Fig.S1 legend).

Due to a similarity in the pathophysiology of sickle cells and RBC from HX-affected people with GoF PIEZO1 variants, and because both types of mutations affect RBC morphology,^30-33^ we first explored if normal and *P. falciparum* infected RBC_E756del_ show any structural abnormalities. As expected,^3^ freshly drawn unprocessed blood from E756del carriers had 6-13% distorted RBC (Fig. 1A). There were frequent erythrocyte ghosts present, suggesting RBC hemolysis *in vivo*, a characteristic of both sickle cell and HX anemias. *P. falciparum* infection of PIEZO1_E756del_ RBC grossly exacerbated erythrocyte cytopathology as also observed in *P. falciparum*-infected sickle cells^34^ but hitherto unreported for PIEZO1_E756del_ RBC. Some infected cells were misshapen, strongly distorted, and severely dehydrated (Fig. 1B). Severe dehydration of iRBC_E756del_ was strikingly like the morphology of dehydrated infected sickle cells.^35^ In addition, some parasites had unusual distortions presenting as semi-split “lobe” shapes, enlarged and dislocated digestive vacuoles (Fig. 1C), and loss of a vacuolar-targeted fluorescent protein in swollen RBC, indicative of a permeable parasitophorous vacuole (Fig. 1D). All these described distortions were more prominent in mature parasites: newly invaded infected cultures had less than 10% distorted infected cells (14 of 149 cells) while more than 50% distorted infected cells (46 of 90 cells) were observed 24 h post infection. The non-infected RBC had a stable fraction of distorted cells in the same cultures (6.5% (24 of 370) cells following parasite invasion and 4.0% (18 of 447) cells 24 h post infection). This distortion of iRBC was stronger on the second day of infection (Fig. 1B-C) consistent with infection induced RBC_E756del_ dehydration. A unique pattern of iRBC_E756del_ dehydration and sickle cells appears to be a common pathophysiological factor we chose to pursue further.

**Fig. 1.**
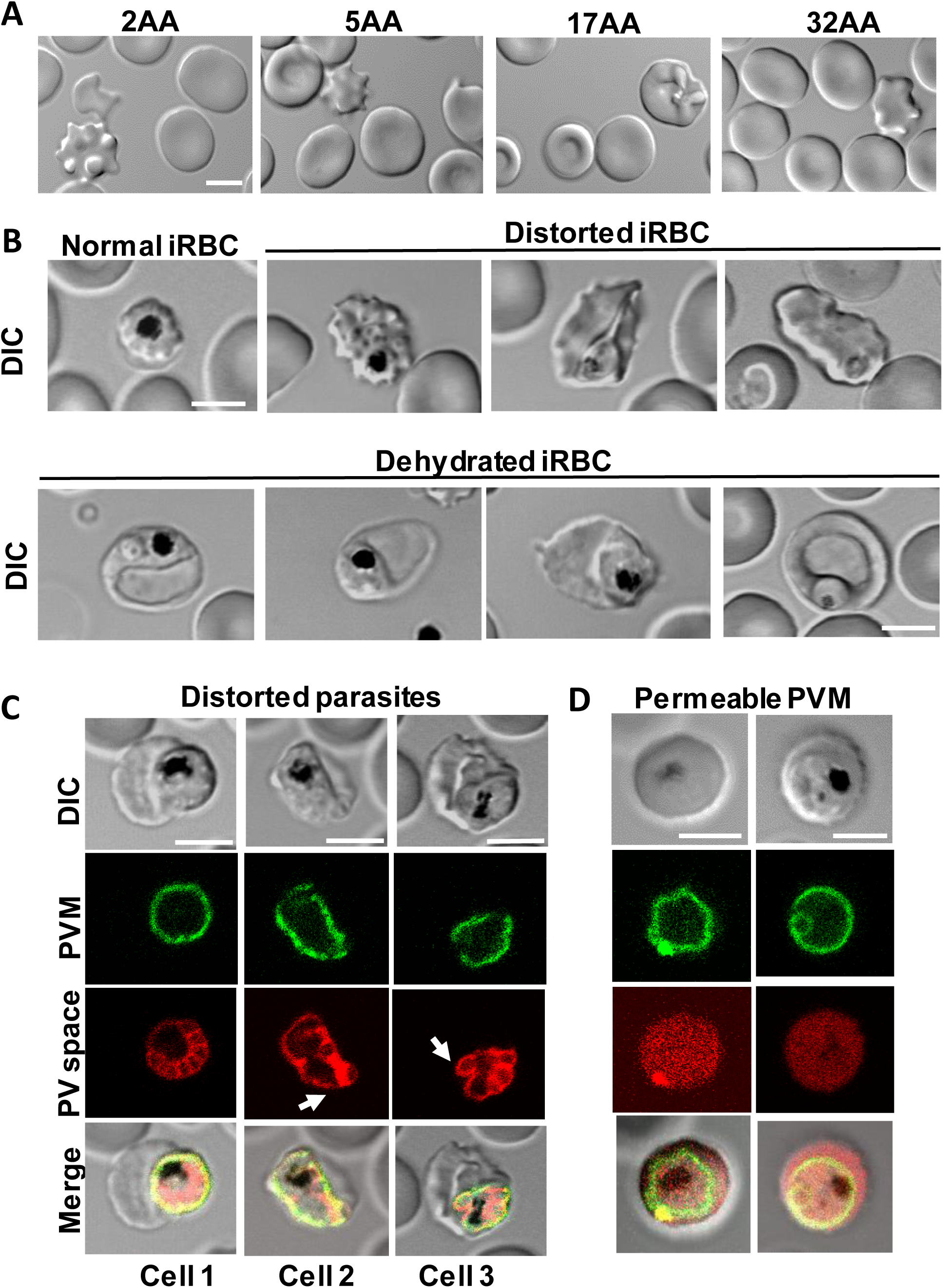
Prominent cytopathology of *P. falciparum* infected RBC_E756del_ and intracellular parasites. A. Optical microscopy images of freshly drawn unprocessed blood from donors 2AA, 5AA, 17AA and 32AA with PIEZO1_E756del_ depict the different types of RBC shape distortions detected in RBC. B. A representative typical iRBC_E756del_ schizont (left image in the upper row) compared with distorted (upper row, three right images) and dehydrated (lower row) iRBC_E756del_. DIC microscopy. C. Morphologically distorted parasites inside iRBC_E756del_ during schizogony: dislocated and enlarged digestive vacuole (cell 1), and semi-split lobe shaped parasites (red arrows, cells 2-3). DIC and fluorescence microscopy. Green color – delineation of the vacuole by the fluorescent vacuolar membrane protein EXP2 tagged with mNeonGreen; red color – parasite-derived, vacuole-containing soluble mRuby3 showing vacuolar space. D. iRBC_E756del_ with the vacuolar membrane permeable to mRuby3. Green and red colors – as described in C. Note RBC swelling. Scale bars in all images = 5 μm.

### *P. falciparum* growth in RBC_E756del_ is cell hydration dependent

To test the hypothesis that RBC_E756del_ in the bloodstream are more dehydrated than in static conditions, due to PIEZO1 activation by shear stress, and may be less suited for parasite replication, the effect of RBC hydration on parasite growth was evaluated first in static cultures with three different RBC hydration levels: 1) the most hydrated RBC with PIEZO1_WT_ from African American and Caucasian donors; 2) moderately dehydrated RBC_E756del_ and 3) the most dehydrated RBC isolated from RBC_E756del_ by Percoll density gradient centrifugation. The C_50_ for each group were 44.35+/-0.69%, mean+/-SEM, n=36 (African American PIEZO1wt donors); 45.61+/-1.11% and mean+/-SEM, n=17 (Caucasian PIEZO1_WT_ donors); 41.10+/-1.16%, mean+/-SEM, n=7 samples from 4 donors (PIEZO1_E756del_ donors); 34.42+/-1.60%, mean+/-SEM, n=5 (dehydrated RBC fraction from PIEZO1_E756del_ donors). Isolated dehydrated RBC_E756del_ from 90% Percoll (density 1.124 g/L) were referred to as the dehydrated cell fraction, or DF (Fig. 2A). They were significantly more dehydrated than WB from the same donors (Fig. 2B) and had more morphologically distorted cells than WB RBC (Fig. 2C). *P. falciparum* growth in WB RBC_WT_ and WB RBC_E756del_ was similar (Fig. 3A-B) but markedly diminished in DF RBC_E756del_ (Fig. 3C). Thus, at some threshold level of RBC_E756del_ dehydration, cells do not support parasite infection.

**Fig. 2.**
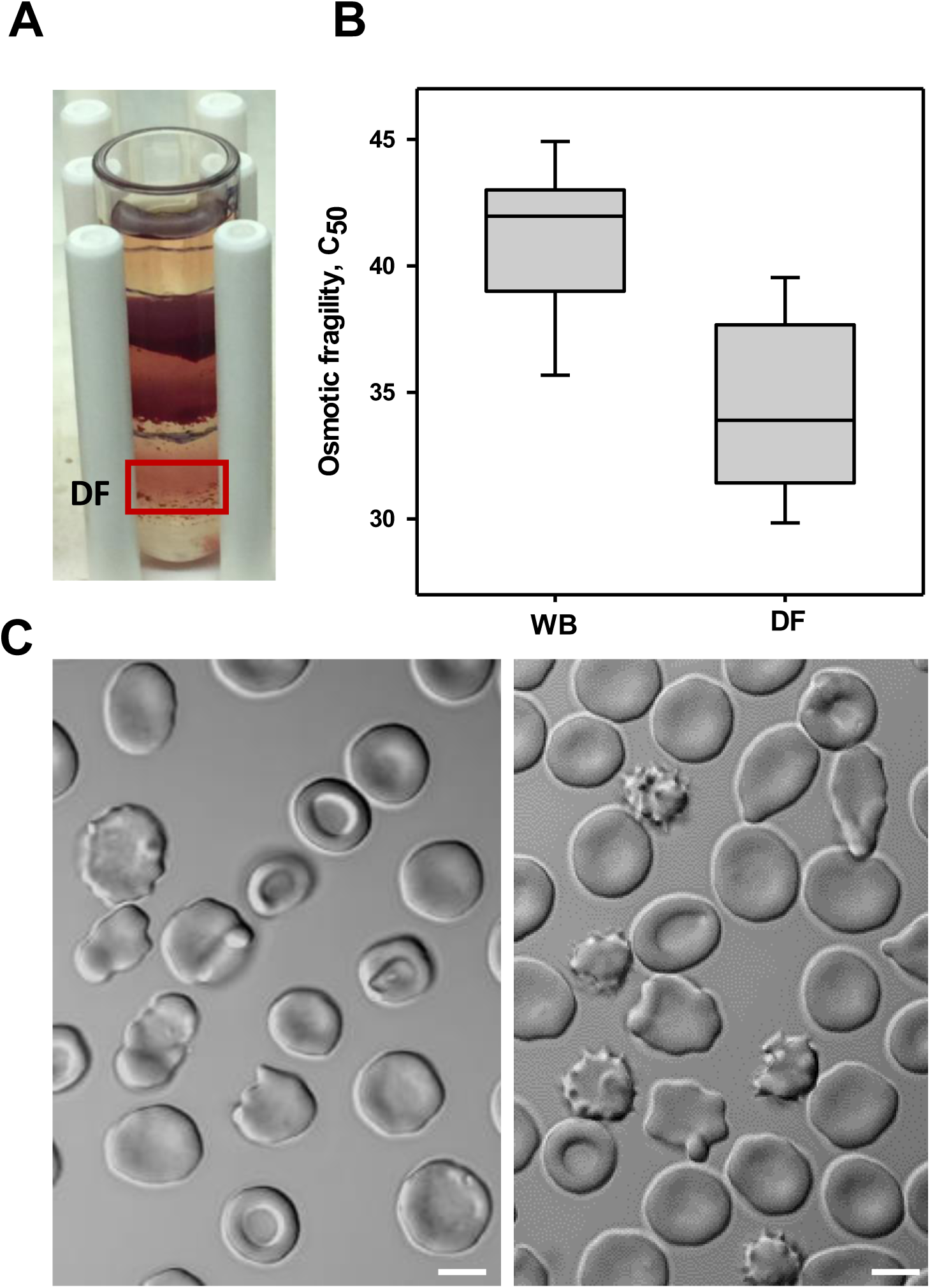
Isolation and characterization of the strongly dehydrated cells from the blood of PIEZO1_E756del_ carriers. A. Steps of 80-90% isotonic Percoll were used for RBC _E756del_ separation based on their hydration status. The bulk of the cells formed a band in the 80% Percoll (upper band), a smaller number of cells were in the 80-90% Percoll interface (middle band), and the smallest number of cells entered the 90% Percoll band (red box). The 90% Percoll fraction was called the “dehydrated cell fraction” or DF. Scale bar = 5 μm. B. RBC hydration, measured in the osmotic fragility assay (C_50_) in whole blood (WB; n=7) and the DF (n=5) of 12 samples from four PIEZO1_E756del_ carriers; cells from the DF were more dehydrated. Bootstrap analysis of the logit transformed C_50_ values (unequal sample size; 10,000 resamples) is consistent with a significant difference in mean hydration. The hypothesis that the two groups have the same mean hydration is rejected at the α = 0.001 level. C. Morphology of RBC from the DF, two representative DIC images. Note multiple crenated and flattened/elongated cells. Scale bars = 5 μm.

**Fig. 3.**
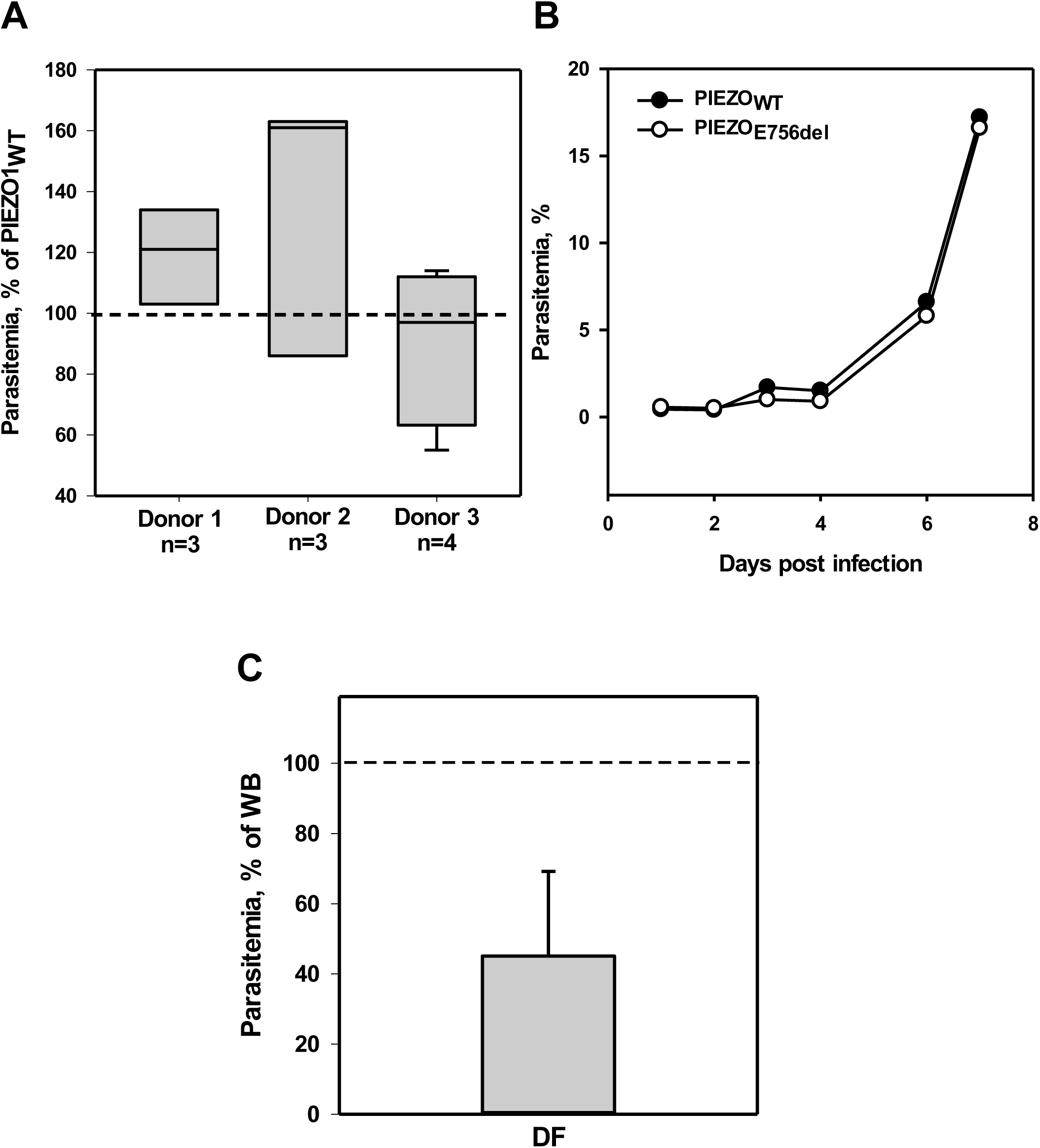
*P. falciparum* growth in RBC_E756del_ is cell hydration dependent. A. Box plots of the parasitemia of the three donors with PIEZO1_E756del_. A one-way ANOVA, of data expressed as a percent of the highest parasitemia in control blood (donors with PIEZO1_WT_), revealed that there was no significant difference in *P. falciparum* parasitemia between the three donors with PIEZO1_E756del_ (F(2,7)=2.07; p=0.20), and, when analyzed as a group, the mean parasitemia was not significantly different from 100%, the control parasitemia in PIEZO1_WT_ (113% +/-24.1%; mean +/-95% CI, n=3-4 for each PIEZO1_E756del/WT_ blood pairs). B. A representative experiment showing similar kinetics of parasite replication in PIEZO1_E756del_ and RBC PIEZO1_WT_ RBC. C. Normalized parasite growth in DF RBC_E756del_ expressed as a fraction of parasitemia in unfractionated RBC from the same donor in side-by-side experiments (blood from 3 donors, 13 independent experiments). Parasitemia was significantly lower than 100% (the upper 99% confidence interval of the log_e_ transformed fractional data is less than l log_e_ (1) = 0; -1.18+/-0.35 (mean+/-SEM); 99% confidence interval = -2.247 and -0.113).

To determine the dehydration-sensitive stage of the *P. falciparum* erythrocyte replicative cycle, parasite invasion, cycle progression, intra-erythrocytic parasite multiplication factor (IMF) and parasite egress were quantitatively assessed in both the DF and WB RBC_E756del_ using optical microscopy.^24-26^ Only parasite invasion of DF cells was significantly inhibited (Fig. 4A-B). Interestingly, distortion with cell crenation did not interfere with parasite invasion (Movie S1), however, only rare invasions of dehydrated elongated RBC were observed (Movie S2), during which time the usual transient cell crenation following parasite invasion was absent;^36^ also see video in Glushakova et al.^37^ Many parasites did not go beyond attachment to the RBC surface (Movie S3). Once inside, the other stages of the parasite erythrocyte cycle were quantitatively like WB RBC_E756del_. The number of merozoites per schizont in DF RBC_E756del_ were not significantly different from WB RBC_E756del_ (IMF in DF RBC = 100.3+/-2.9% of IMF in WB, mean +/-SEM, n=3) (Fig. 4C, a representative experiment). The end of cycle time and kinetics of egress in DF cultures were not significantly different from WB (DF =100.4+/-0.8% of those in WB RBC, mean+/-SEM, n=3) (Fig. 4D, a representative experiment). Egressed parasites were fully separated (Fig. 4E). Thus, crenation of RBC _E756del_ was compatible with infection initiation but distortions that stretched the RBC restricted parasite invasion. Parasite egress from the dehydrated iRBC_E756del_ showed prominent deviations, reminiscent of those in dehydrated sickle cells:^35^ delayed RBC membrane rupture and vesiculation of the RBC membrane prior to membrane opening (Movie S4). Thus, parasite infection of RBC_E756del_ in the absence of PIEZO1 activation was unimpaired in hydrated RBC but was hampered in dehydrated cells due to limited parasite invasion of and defective parasite egress from these cells.

**Fig. 4.**
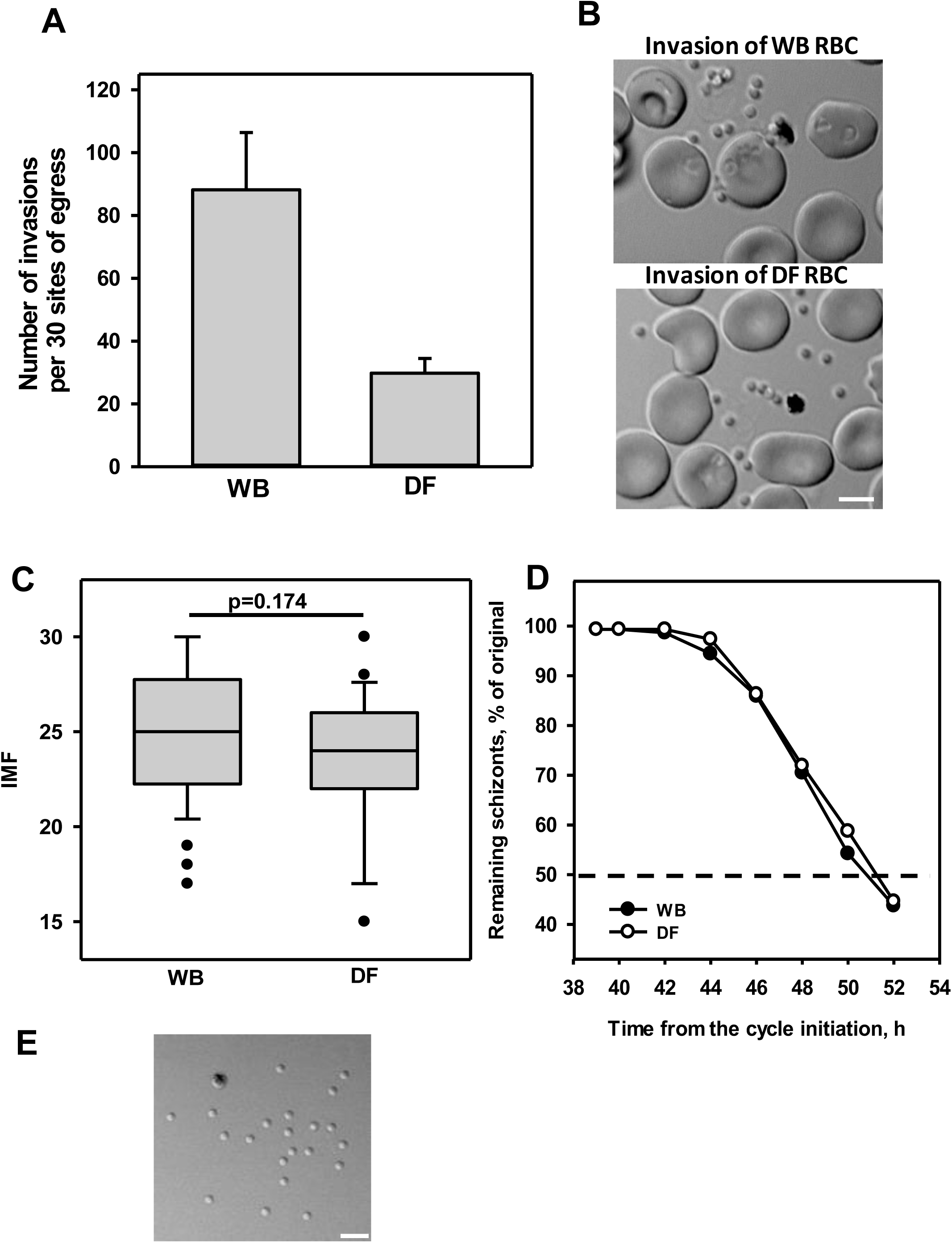
Inefficient parasite growth in DF RBC_E756del_ in comparison with unfractionated RBC from the same donors correlates with the defect in parasite invasion of dehydrated RBC. A. Deficiency in parasite invasion of dehydrated cells correlates with inefficient parasite replication in these cells (blood from 3 donors, 5 independent experiments). Number of invasions was evaluated in 30 sites of parasite egress for both WB and DF of donor RBC per each experiment; parasite invasion of dehydrated RBC (WB vs. DF invasions; 88.2+/-18.2 vs 29.8+/-4.7, mean+/-SEM, n=5, p=0.03, unequal variance t-test). B. The representative sites of parasite egress and invasions in experiments with WB (upper image, 8 invasions in the neighboring RBC) and DF (lower image, 2 invasions in the neighboring normal RBC) in side-by-side experiments with the blood from the same PIEZO_E756del_ donor. DIC microscopy. C. IMF evaluation (i.e., the number of parasites egressed from one schizont; data plotted as median and percentiles, 30 sites of egress were evaluated for each type of RBC, as illustrated in panel E), a representative experiment. D. Parasite egress kinetics to compare the length of replicative cycle in WB and DF experiments; a representative experiment. E. Inspection of merozoite formation and their separation upon egress, DIC microscopy. Scale bar in all images = 5 mm.

### PIEZO1 agonist Yoda1 inhibits parasite growth in RBC_E756del_ and induces cell hemolysis

Next, PIEZO1 was chemically activated using the PIEZO1 agonist Yoda1, thus increasing RBC dehydration in the entire RBC_E756del_ population *in vitro* at the time of parasite invasion.^16^ We confirmed that low concentration of Ca^2+^ in complete medium is sufficient to initiate cell volume loss, expressed as appearance of echinocytes, or crenated cells, due to activation of PIEZO1 and Gardos channels upon Yoda1 addition (Fig. 5A, upper image, non-infected RBC, red arrow). Cell distortion was only partially reversed by Yoda1 removal: approximately half of the cells were distorted 6 days after treatment (Fig 5A, middle image). Intriguingly, iRBC_E756del_ show no RBC crenation but show cell flattening and stretching, typical for dehydration (Fig. 5A, upper (white arrow), and lower images; for comparison with dehydrated infected sickle cells see Glushakova et al.^35^ Both 5 and 10 μM Yoda1 treatments for 1 h during infection initiation inhibited parasite replication in WB and DF static cultures (Fig. 5B). The appearance of erythrocyte ghosts in the Yoda1-treated cultures (Fig. 5B, inset, red arrow) prompted evaluation of RBC hemolysis in these experiments. Cumulative hemolysis over the entire experimental time span showed that Yoda1 induced hemolysis in cultures with both significant dose and cell hydration factors (Fig. 5C). Thus, dehydration of cells, following Yoda1 activation of PIEZO1, inhibits parasite replication in RBC_E756del_ and induces the additional pathophysiological effect on erythrocytes, i.e. hydration-dependent cell hemolysis.

**Fig. 5.**
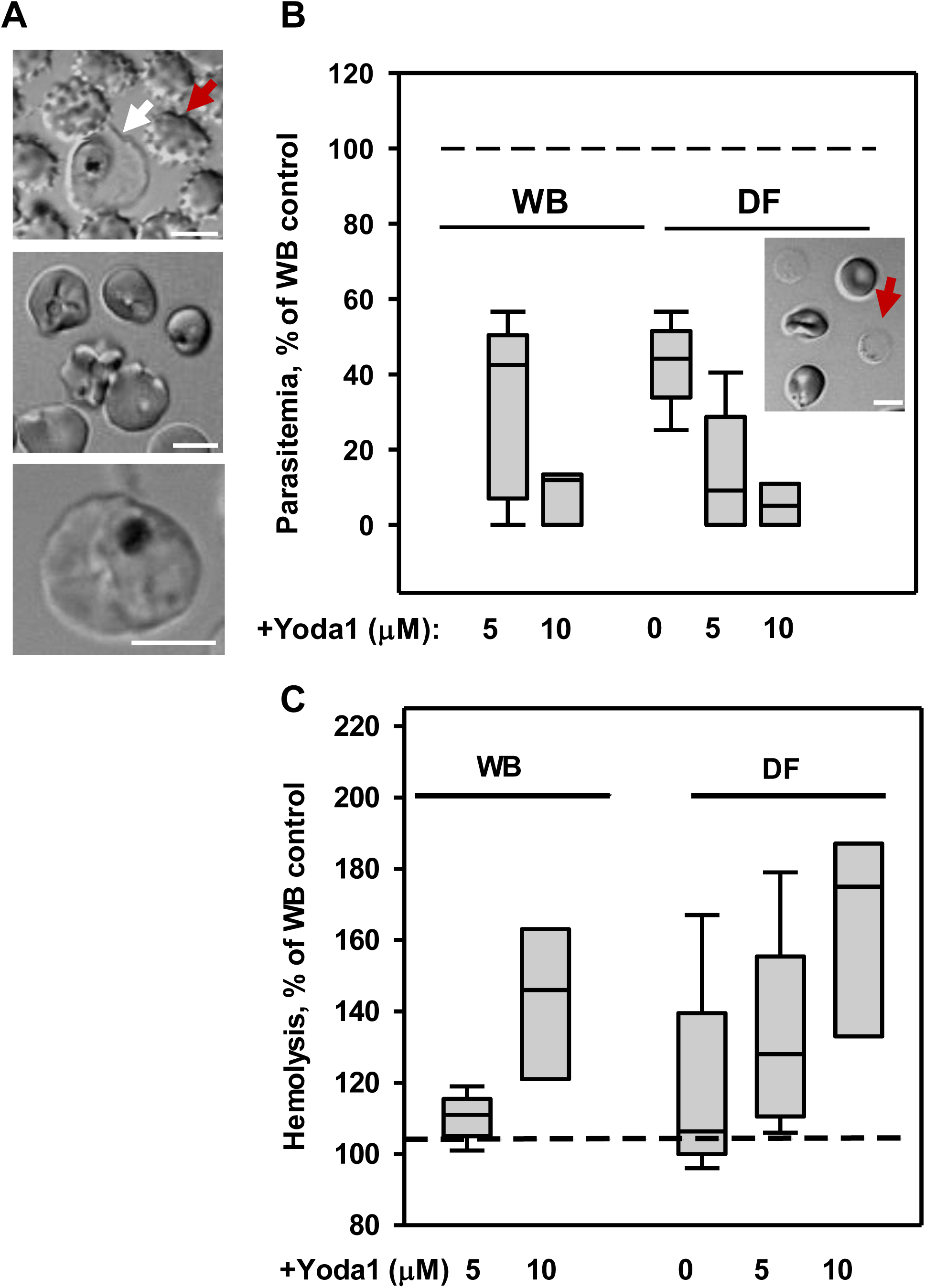
PIEZO1 agonist Yoda1 negatively affects RBC_E756del_ morphology, parasite replication and RBC _E756del_ stability. A. Echinocytosis (crenation) of normal RBC (red arrow) and stretching/flattening of infected cell (white arrow) within a few minutes post Yoda1 exposure (upper image); distorted morphology of Yoda1-treated RBC lasts for days after drug removal (middle image); dehydrated iRBC on day 6 of the experiment (lower image), DIC microscopy. Scale bar in all images = 5 μm. B. Strong inhibition of parasite replication in WB and DF RBC, treated with Yoda1 for 1h in static cultures (two donors, 5 independent experiments: data plotted as median and percentiles). To evaluate the data, the 0% values were first adjusted to 0.05% based on the random sampling of 1,500 cells, data was logit transformed, and a 2-factor ANCOVA analysis was performed using Yoda1 dose as a continuous variable. With no evidence for a significant interaction term, a model with no interactions was used. Yoda1 treatment was significantly inhibitory for parasite growth in the tested concentrations (F(1,18)=5.35, p=0.033); cell hydration was not a significant factor for this data set (F_hydration_(1,18)=0.6, p=0.447). Inset: RBC ghosts (red arrow) in the Yoda1-treated cultures. Scale bar = 5 μm. C. Yoda1 induces dose-dependent and cell-hydration dependent erythrocyte hemolysis in the static cultures of infected RBC. Hemolysis was assessed by detection of hemoglobin (Hb) release into the culture medium during the entire experiment (cumulative hemolysis, Hb adsorption at 540 nm) and normalized per hemolysis in the control WB culture (2 donors, 5 experiments; data plotted as median and percentiles). A 2-factor ANCOVA analysis of the log_e_ transformed fold change, with dose as a continuous variable, indicate a significant linear dependence on the Yoda1 dose and blood hydration (F_yoda1_(1,18)=12.33, p = 0.002 and F_hydration_(1,18=4.7, p=0.044).

### Activation of PIEZO1 E756del by shear stress induces cell hemolysis and has a strong inhibitory effect on parasite replication in RBC_E756del_

To mimic blood stream turbulence that creates shear stress activation of PIEZO1,^38-39^ experiments were performed with samples placed on a “belly-button” rocker/shaker platform (see the Method section). Both WB and DF RBC_E756del_ were tested side by side in the static and rocking (shear stress) conditions. Shear stress significantly inhibits parasite replication within the first replicative cycle (see representative kinetics, Fig. 6A-B). The growth of parasitemia was abolished after infection initiation or sharply declined if shear stress was applied in the middle of the experiment (Fig.6C). In the latter case, the inhibition of parasite replication was 79.4% and 87.3% for the first two days in comparison with the static cultures (two independent experiments). Shear stress led to swelling and permeabilization of iRBC (inset in Fig. 6A). Like PIEZO1_E756del_ activation with Yoda1, shear stress activation of the channel led to RBC_E756del_ hemolysis (Fig 6D-E). Hemolysis was more prominent in DF than in WB cultures. RBC from two PIEZO1_E756del_ donors had different sensitivity to shear stress (compare Fig. 6D and 6E); one donor’s blood showed cell hemolysis even under static conditions (not reflected in this graph due to the data normalization used) and a much higher hemolysis under shear stress conditions (Fig. 6E). On day 3 of the experiment, fluorescence microscopy of cells labeled with the membrane dye FM64 revealed RBC ghosts, fragmented cell membranes and hemolyzed infected cells (Fig. 6F, red color). The physical instability of the RBC_E756del_ under shear stress supports the hypothesis that these cells are not competent for supporting parasite infection. Thus, activation of PIEZO1_E756del_ channel, *in vitro*, by the channel agonist Yoda1 and by continuous turbulent shear stress inhibits parasite infection by inducing cell hemolysis.

**Fig. 6.**
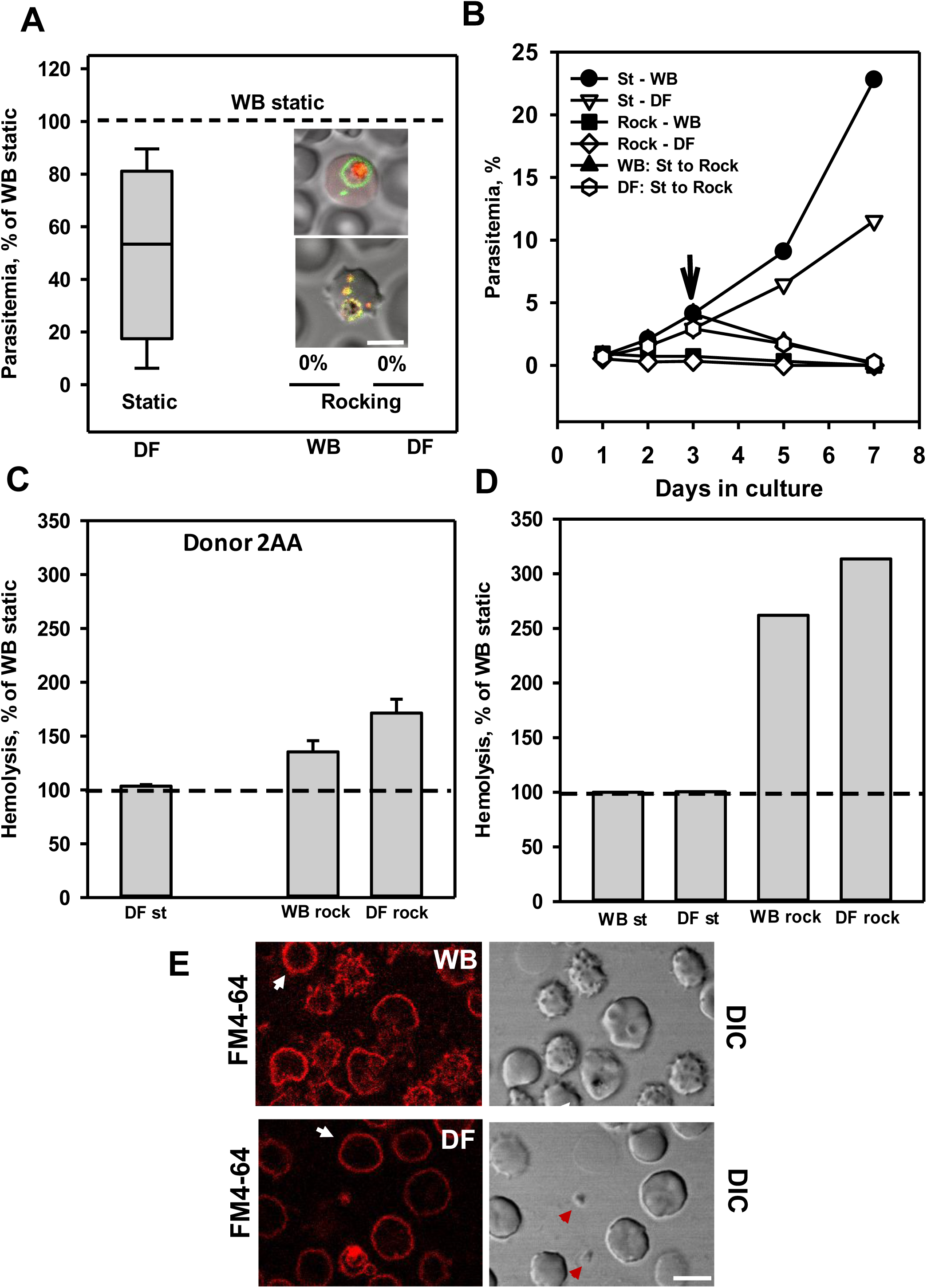
Shear stress inhibits parasite replication and leads to cell hemolysis in the RBC_E756del_ cultures. A. Shear stress blocks parasite replication. The highest achieved parasitemia in the WB static culture was used as a normalizing value for presentation of parasite replication in all cultures (two different donors, 4 experiments, data plotted as median and percentiles). Two-factor ANOVA analysis after correction for “0” percentage and log_e_ transformation indicate that there is a significant dependence on the culture conditions (static vs. rocking, F_Shear-Stress_(1,12)=23.0, p=0.0004). Inset: iRBC swelling and membrane permeabilization in rocking condition. Color code as in Fig. 1C. Fluorescence and DIC microcopy. Scale bar = 5 μm. B. A representative experiment shows the kinetics of parasite replication in the static (St) and rocking (Rock) conditions and a sharp decline in parasite replication upon transfer of the static culture to the rocking platform; arrow shows the time of cultures transfer. C. Shear stress induces both cell-hydration dependent (two-factor ANOVA, F_Hydration_(1,12)=6.62, p=0.02) and culture-condition dependent (two-factor ANOVA, F_Shear-Stress_(1,12)=28.55, p=0.0002) RBC hemolysis (one donor 2AA, three blood donations, 5 independent experiments, mean +/-SEM). D. Blood from the donor 32AA shows much stronger cell hemolysis in the rocking cultures than blood of 2AA donor, as shown in (C), one experiment. E. Fluorescence and DIC microscopy images of RBC after 3 days in the shear stress culture from donor 32AA confirms physical cell damage. Note pre-hemolytic DF cell phenotype, i.e., rounded swelled shape. Red color: FM4-64 membrane dye showing erythrocyte ghosts (white arrow) and membrane fragments (small red dots, red arrowhead) from the hemolyzed RBC. Scale bar = 5 μm.

## Discussion

Here, we demonstrate a connection between increased RBC_E756del_ dehydration, following activation of GOF PIEZO1, with increased sensitivity to hemolysis and impaired *P. falciparum* replication. Even in static culture, with presumably inactive PIEZO1_E756del_, *P. falciparum* replication was diminished in isolated dehydrated cells. The inverse correlation between RBC dehydration and parasite replication in isolated DF cells can resolve a discrepancy in the results on the effect of PIEZO1_E756del_ mutation on parasite replication *in vitro*^3,7^ where a small number of donors with different mean RBC dehydration was used. In our study, RBC_E756del_ showed unimpaired parasite growth, in vitro, in side-by-side experiments with RBC_WT_. Thus, sufficient RBC_E756del_ hydration, when PIEZO1 is inactive, makes these cells suitable for *P. falciparum* replication. However, in static cultures, some infected cells displayed an unusually strong dehydration phenotype like those observed in infected sickle cells.^35^ This morphological similarity may be explained by similar mechanisms of cell dehydration, i.e., over-activated PIEZO1 and Gardos channels in combination with overwhelmed Ca^2+^ pumps.^18,20,30,40-42^ RBC dehydration appears to be exacerbated by infection in both cell types leading to aberrant parasite egress from a sub-population of cells (this and two other studies).^34,35^ Most importantly, dehydrated naïve RBC_E756del_ did not allow efficient invasion by *P. falciparum*, severely limiting parasite replication in cultures of dehydrated cells. Together with the early data on inefficient parasite replication in dehydrated HX RBC of unknown etiology^27^ and the recently revealed negative correlation between RBC dehydration alleles and parasite fitness^11^, our results confirm an inverse relationship between RBC dehydration and parasite replication in RBC. In addition, RBC dehydration may increase membrane tension that negatively affects parasite invasion of RBC, as described for the Dantu blood group.^43^ Thus, impaired parasite invasion of, and defective parasite egress from dehydrated cells are two defects in the parasite replicative cycle that occur in RBC_E756del_ in the static cultures, without PIEZO1 activation. If during blood circulation *in vivo* cell dehydration ensues due to PIEZO1 activation, then these replication defects can contribute to a milder disease.^3,7^

We recapitulated this scenario *in vitro* by activating PIEZO1_E756del_^16,20^ and discovered an additional phenomenon: dehydration-dependent cell hemolysis under conditions that showed severely diminished parasite replication in RBC_E756del_. The mechanism of cell hemolysis upon PIEZO1 activation was not addressed in our study. As discussed in the introduction, the E756del mutation reduces PIEZO1 inactivation, leading to higher cytoplasmic [Ca^2+_free_^], which could activate calpain and thus increase cell fragility due to digestion of the RBC cytoskeleton. Alternatively, the longer open state of PIEZO1_E756del_, as a non-specific cation channel PIEZO1, would directly conduct Na^+^ to enter down its activity gradient, which combined with the slow chloride permeability of RBC could be sufficient to promote colloid hemolysis despite KCNN4 activation. Whether parasite infection increases PIEZO1_E756del_ activation and further dehydrates the host cell, or growing parasites simply make dehydrated RBC_E756del_ less deformable and more sensitive to shear stress damage^44^ or both, remains to be established. Irrespectively, the consequences for a parasite attempting to replicate in dehydrated cells, could be shear stress-induced RBC hemolysis that prematurely terminates the replicative life cycle. Experimentally confirmed, dehydration-dependent hemolysis of RBC highlights another commonality between RBC_E756del_ and sickle cells: both pathologies generate dehydrated cells, and both are characterized by RBC hemolysis.^31,45^

The correlation between RBC dehydration and hemolysis in PIEZO_E756del_ activated RBC suggests the following mechanism of protection of PIEZO1_E756del_ carriers from severe forms of malaria: gain of function PIEZO1_E756del_ becomes activated by shear stress and leads to increased RBC dehydration that, in turn, leads to increased RBC hemolysis and inefficient parasite replication. More generally, cell dehydration could emerge as a factor associated with the amelioration of malaria both in malaria patients and in rodent malaria models due to i) sickle hemoglobin,^30,31,45^ ii) deficiency in PMCA4 production,^46^ iii) gain of function KCC1 co-transporter^47-48^ and iv) gain of function PIEZO1_E756del_.^3,7^ Further investigations into the protective mechanisms of erythrocyte dehydration and erythrocyte membrane pathology on malaria parasite biology and malaria pathogenesis is warranted.

## Supporting information

See manuscript file

See manuscript file

See manuscript file

See manuscript file

## Acknowledgements

We thank all the anonymous donors of blood who contributed to the results of this study. We thank Dr. Josh Beck for providing a strain of *P. falciparum* expressing the fluorescently tagged proteins, Drs. Kamran Melikov and Matthias Garten for consultation in microscopy. This study was funded by the Intramural Research Program of the *Eunice Kennedy Shriver* National Institute of Child Health and Human Development.

## Authorship

S.G., H.W, Y.K. and J.Z. conceived and designed experiments. S.G., L.B. and H.W. performed experiments. P.S.B. performed statistical analysis. All authors analyzed data and wrote the manuscript.

## Supplemental data to the manuscript

### Supplemental Fig. 1 legend

**Fig. S1.**
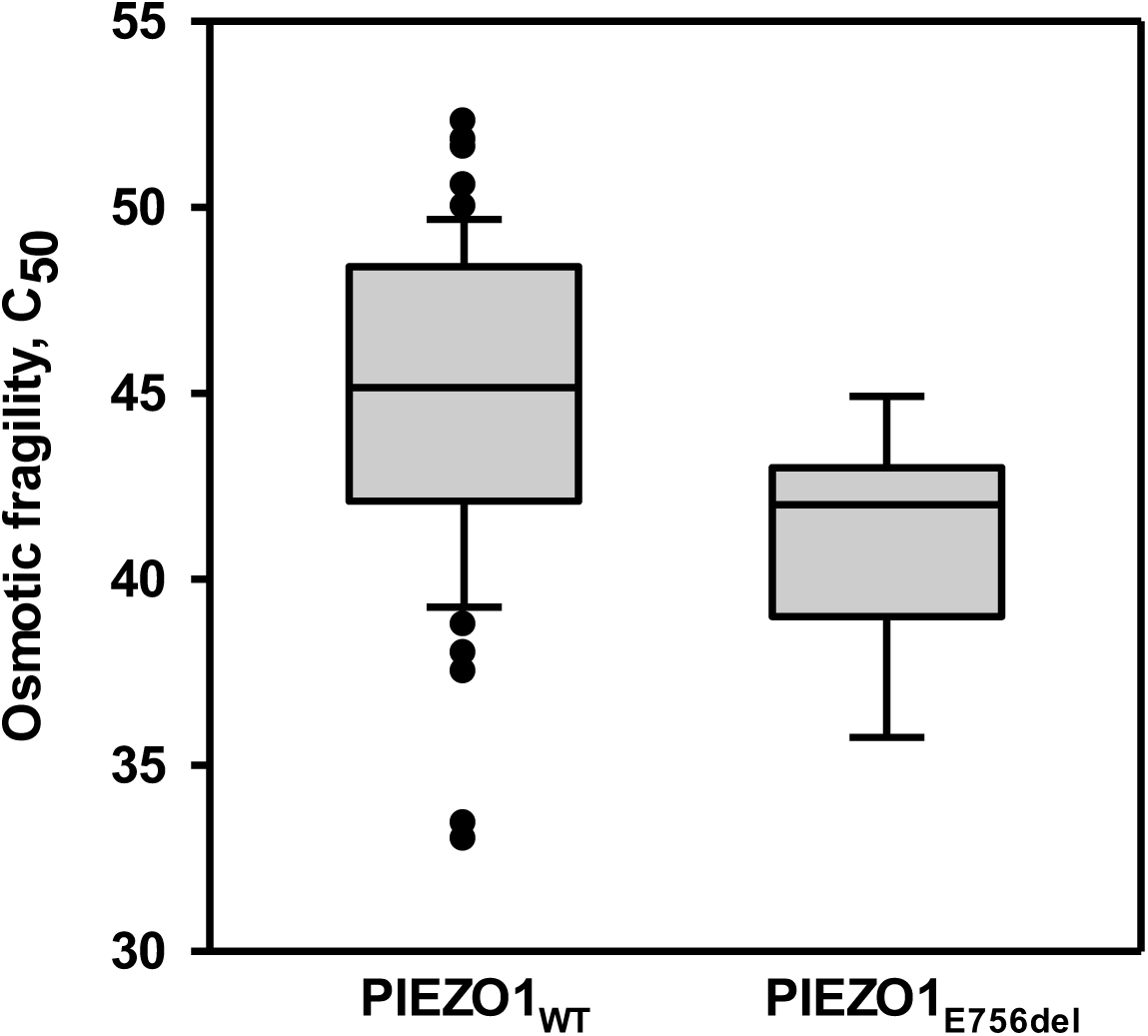
GoF PIEZO1 _E756del_ mutation makes RBC less hydrated than PIEZO1_WT_ under static *in vitro* conditions with presumably inactive PIEZO1 channel. N=7 for PIEZO1 GoF blood (African American donors) and n=53 for PIEZO1_WT_ blood (African American and Caucasian donors). To evaluate the difference between cell hydration in RBCs with PIEZO1_WT_ and PIEZO1_E756del_, data from all PIEZO1_WT_ donors (n=53) were compared with data from PIEZO1_E756del_ donors (n=7). A non-distributional bootstrap analysis of the logit transformed C_50_ values (unequal sample size; 10,000 resamples) was used. The null hypothesis that the two groups have the same mean hydration is rejected at the α = 0.001 level.

### Supplemental Movie legends

**Movie 1: *P. falciparum* can invade distorted RBC**_**E756del**._

Parasite egress and invasion of two neighboring erythrocytes. Four total invasions: three in the left RBC and one in the upper distorted RBC. An invasion induces a mild echinocytosis of the distorted RBC in comparison with a strong echinocytosis of the biconcave RBC. DIC microscopy. Note, that intervals between the frames in this and the following movies are variable for minimization of cell photodamage.

**Movie 2: Parasite invasion of an elongated dehydrated RBC**_**E756del**_ **is not followed by post-invasion RBC echinocytosis**.

Parasite egress and invasion of an elongated RBC has minimal if any post-invasion RBC echinocytosis. DIC microscopy.

**Movie 3: A failed attempt of parasite invasion of elongated dehydrated RBC**_**E756del**_ **despite parasite attachment to the cell surface**.

Parasite egress from the dehydrated infected RBC_E756del_ and invasion of three biconcave neighboring RBC followed by strong host cell echinocytosis. The elongated dehydrated RBC was not invaded by the parasite despite its attachment to the cell surface. DIC microscopy.

**Movie 4: Distortion of parasite egress from the infected RBC**_**E756del**_ **featured a delayed rupture of vesiculated host cell membrane**.

Depiction of parasite egress from an infected RBC_E756del_ that has a long time (21 min vs. the average 7 min interval; see Glushakova et al., 2018)^28^ between vacuolar membrane and RBC membrane rupture leading to parasite egress and dispersion. Unusual RBC membrane vesiculation initiated ∼ 9 min before parasite egress. Both abnormalities were often observed in infected sickle cells (see Glushakova et al., 2010).^35^ Fluorescence and DIC microscopy. Green color – delineation of the PVM containing membrane protein EXP2-mNeonGreen. Red color – soluble vacuolar protein mRuby3, showing the vacuolar space.

